# How to test for phasic modulation of neural and behavioural responses

**DOI:** 10.1101/517243

**Authors:** Benedikt Zoefel, Matthew H Davis, Giancarlo Valente, Lars Riecke

**Author notes:** Corresponding author: Benedikt Zoefel MRC Cognition and Brain Sciences Unit University of Cambridge 15 Chaucer Rd Cambridge CB2 7EF United Kingdom.

## Abstract

Research on whether perception or other processes depend on the phase of neural oscillations is rapidly gaining popularity. However, it is unknown which methods are optimally suited to evaluate the hypothesized phase effect. Using a simulation approach, we here test the ability of different methods to detect such an effect on dichotomous (e.g., “hit” vs “miss”) and continuous (e.g., scalp potentials) response variables. We manipulated parameters that characterise the phase effect or define the experimental approach to test for this effect. For each parameter combination and response variable, we identified an optimal method. We found that methods regressing single-trial responses on circular (sine and cosine) predictors perform best for all of the simulated parameters, regardless of the nature of the response variable (dichotomous or continuous). In sum, our study lays a foundation for optimized experimental designs and analyses in future studies investigating the role of phase for neural and behavioural responses. We provide MATLAB code for the statistical methods tested.

## 1. Introduction

Neural oscillations are cyclic variations in the excitability of neuronal ensembles. Oscillatory phase indexes the instantaneous state of excitability (Buzsáki and Draguhn, 2004) and hence correlates with neuronal firing in intracranial recordings (Kayser et al., 2015). The phase of neural oscillations estimated using non-invasive methods such as electro- or magnetoencephalography (EEG/MEG) has been shown to influence human perception and other aspects of cognition (shown schematically in Fig. 1A). For instance, in both the visual and somatosensory systems, the detection of a near-threshold stimulus and the ability to distinguish two rapidly presented stimuli correlate with EEG/MEG phase (Ai and Ro, 2014; Baumgarten et al., 2015; Busch et al., 2009; Mathewson et al., 2009; Milton and Pleydell-Pearce, 2016; Ronconi et al., 2017; but see Ruzzoli et al., 2019). In these studies, stimuli are often presented at an a-priori unknown, random neural phase in each trial; this leads to the possibility of testing how neural or behavioural responses relate to the phase of spontaneous (i.e. stimulus-unrelated) oscillations extracted post-hoc from concurrent EEG/MEG recordings.

**Figure 1.**
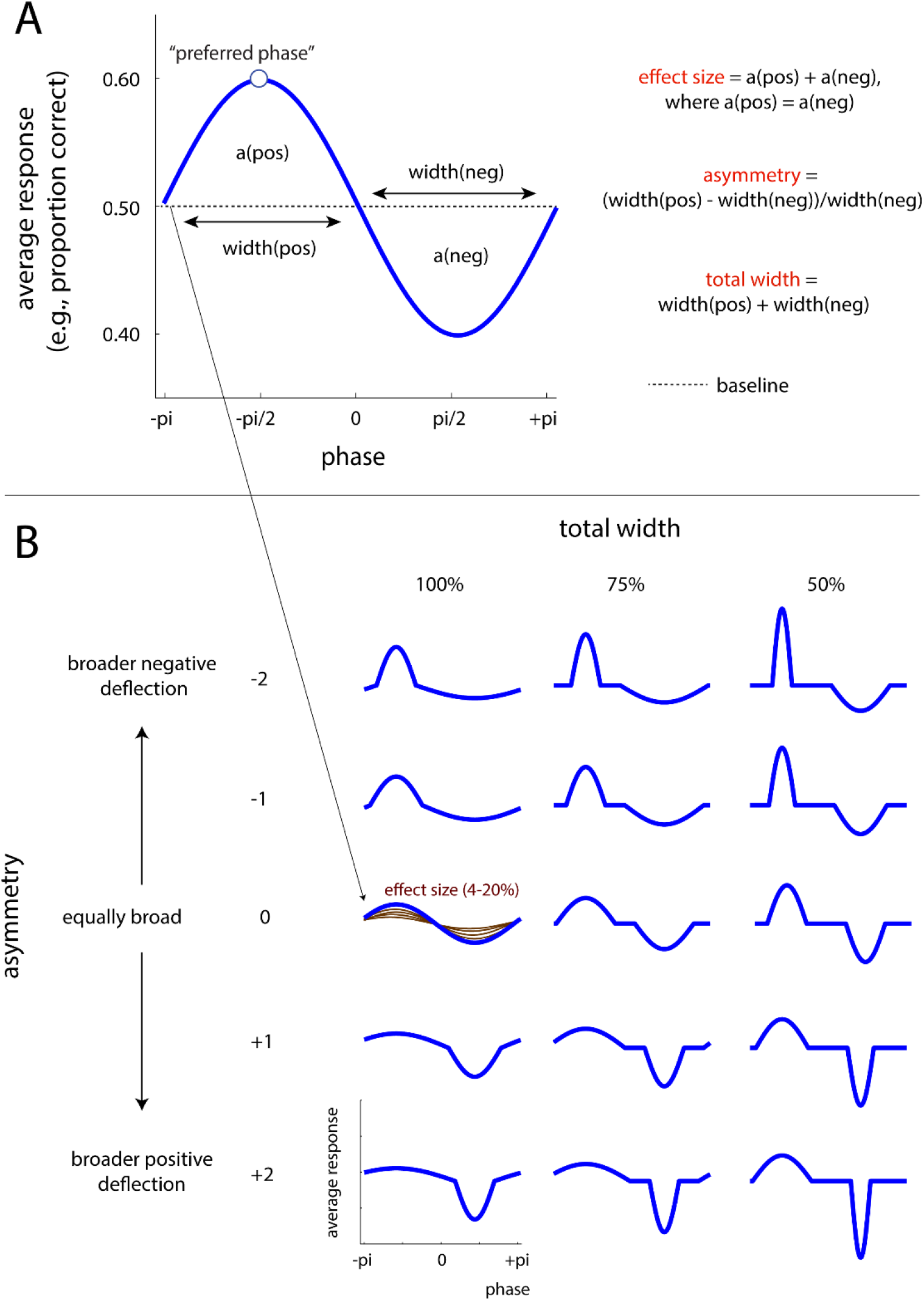
Modelling the phase effect. **A**. Definition of neural parameters. The vertical axis represents the average response (e.g., proportion of hits for a dichotomous response variable, average scalp potential for a continuous response variable) at each simulated phase. This corresponds to *model(phase)* in Section 2.1.1. **B**. Simulated values for neural parameters. These parameters produce different effect shapes that can be sinusoidal (*total width =* 100%, *asymmetry =* 0) or not. Note that *effect size* is identical for all of the blue curves shown. For one exemplary combination of asymmetry and total width, all possible *effect sizes* are shown in brown.

An alternative paradigm exploits the observation that neural oscillations align to rhythmic stimulation. In addition to sensory input (Henry and Obleser, 2012; Hickok et al., 2015; Keitel et al., 2017; Mathewson et al., 2012; Spaak et al., 2014), a commonly used stimulus for this purpose is transcranial alternating current stimulation (tACS; Herrmann et al., 2013; Riecke and Zoefel, 2018; Zoefel and Davis, 2017). In such studies, the phase of the electric stimulus serves as a surrogate for neural phase, and the effect of this presumed neural phase on neural or behavioural responses can be assessed. Imposing phase externally in an experimentally controlled manner allows testing for causal effects of phase, but often limits the number of discrete phase values that can be tested.

Irrespective of the exact paradigm used, a question common to all the studies described above is whether and how neural or behavioural responses are influenced by phase. Importantly, there is currently no consensus on how such phase effects can be best revealed statistically (cf. Asamoah et al., 2019). One common statistical approach is to compare responses across trials grouped into phase bins using *parametric tests* (Baumgarten et al., 2015; Busch et al., 2009; Busch and VanRullen, 2010; Chakravarthi and VanRullen, 2012; Gundlach et al., 2016; Madsen et al., 2019; Mathewson et al., 2009; Neuling et al., 2012; Ng et al., 2012; Riecke et al., 2015a, 2015b, 2018; Ruzzoli et al., 2019; Wilsch et al., 2018; Zoefel et al., 2018a; Zoefel and Heil, 2013). However, individuals are often found to differ in their *“preferred” phase* (defined as the phase observed to yield the participant’s maximum response; see Fig. 1A and 2A; e.g., Busch et al., 2009; Riecke et al., 2018). Consequently, data is commonly reported from phase bins that are re-aligned relative to the preferred phase for each participant. This alignment makes it mandatory to exclude the phase bin used for alignment, which therefore reduces potentially important phase-related variance from subsequent analyses of the aligned data.

**Figure 2.**
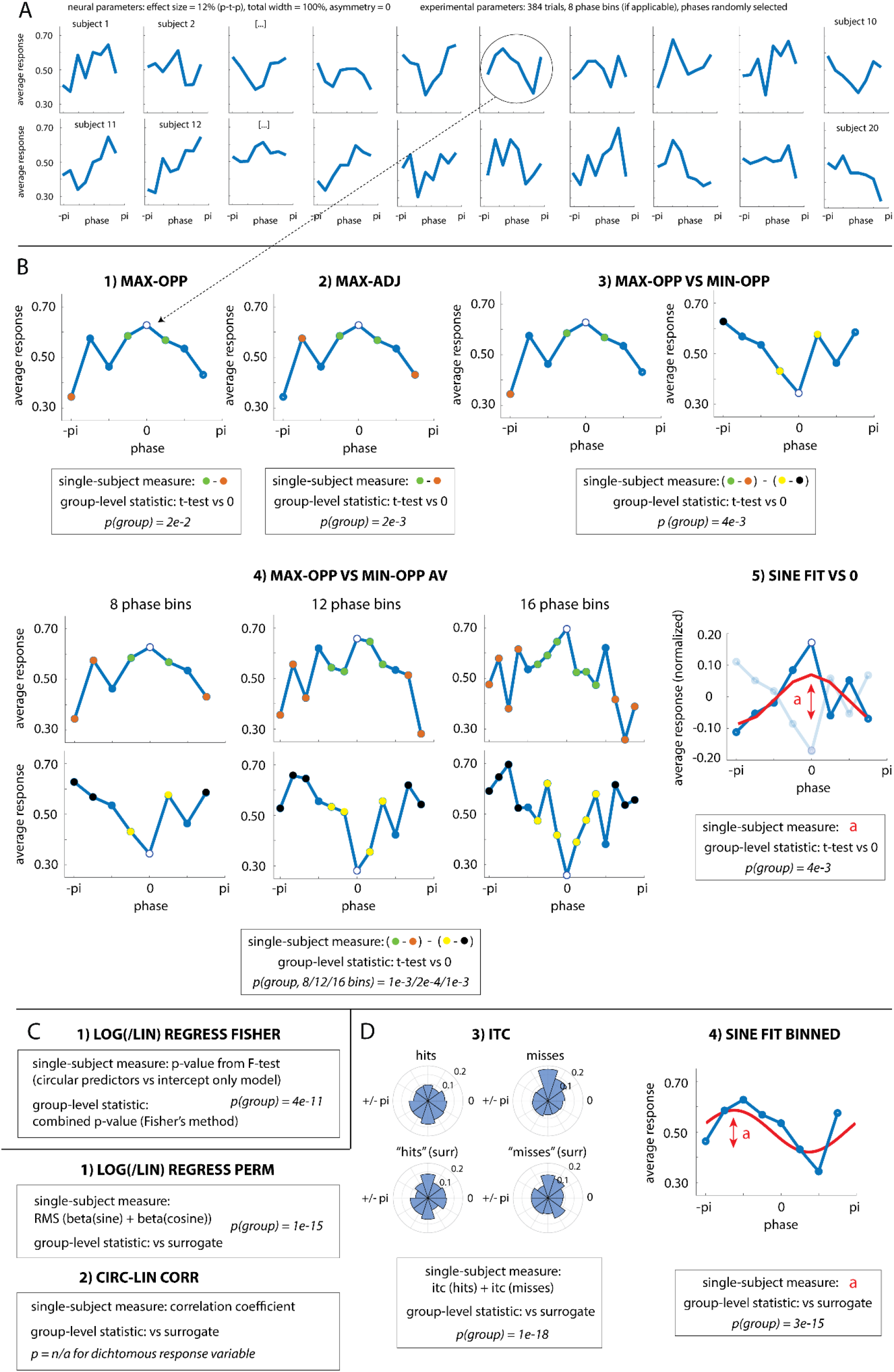
Illustration of statistical methods. **A**. Data from 20 virtual participants in one experiment for a given combination of (neural and experimental) parameters and a dichotomous response variable. Note that the preferred phase (the phase yielding the maximum response) differs across participants. **B-D**. Statistical methods that were included in the present study, divided into three categories. For all methods, the single-subject measure of the hypothesized phase effect is visualized and/or described based on data from one exemplary subject. If methods divide data into phase bins, these are shown with circles. For alignment-based methods (B), the bin used for alignment is shown as an open circle. For all methods, p-values shown were obtained for the group level, by applying the respective method to the data shown in A. The panel illustrating method ITC (D, 3) depicts phase distributions for hit and miss trials (observed data and surrogate distribution). See Section 2.2 for further details.

A second common statistical approach is the use of non-parametric, *permutation-based methods*: Here, the observed data is compared with a surrogate distribution, designed to simulate outcomes due to random fluctuations in the data (i.e. the null distribution). This distribution can be constructed by repeatedly shuffling responses (or phases) associated with single trials in the original dataset and re-running the original analysis on the permuted datasets. Most studies have quantified phase effects by comparing phase distributions of trials that were grouped according to a specific observed behavioural response (e.g., “hit” vs “miss”) (Busch et al., 2009; Dugué et al., 2011; Hanslmayr et al., 2013; Harris et al., 2018; Ng et al., 2012; Ronconi et al., 2017; Ruzzoli et al., 2019; Strauß et al., 2015; ten Oever and Sack, 2015; Wutz et al., 2016). Other studies have used regression-based methods (Kayser, 2019; Kayser et al., 2016; McNair et al., 2019) or circular-linear correlations (Busch and VanRullen, 2010; Chakravarthi and VanRullen, 2012) in combination with permutation tests.

In this study, we conducted Monte-Carlo simulations to determine how reliably different statistical methods can detect a phasic modulation of neural or behavioural responses. In principle, these methods can be applied to reveal phase effects in any type of signal, including but not restricted to oscillatory ones. Indeed, some species show phasic modulation of neural activity in the absence of neural oscillations (Eliav et al., 2018). Nevertheless, we here implicitly assumed that the simulated phases stem from an oscillatory (i.e. narrow-band) signal whose phase can be readily interpreted. Methods to detect specifically oscillatory activity have been described elsewhere (e.g., Haller et al., 2018; Watrous et al., 2018; Zoefel et al., 2018b).

We tested how two sets of parameters impact the sensitivity of the tested methods: Parameters that describe the nature of the phase effect and reflect underlying neural processes (Fig. 1), henceforth termed *neural parameters*, and parameters that depend on the specific experimental design, termed *experimental parameters* (see corresponding sections in Material and Methods). We contrasted the two categories of statistical methods described above (*parametric alignment-based* vs *permutation-based*; Fig. 2B,D), and an additional category of parametric regression-based methods requiring no alignment (Fig. 2C). We investigated phase effects on two different types of response variables: dichotomous (“hit” vs “miss”) and continuous (e.g., scalp potentials or response times). Together, the results of our simulations serve to guide researchers in choosing the most appropriate method to detect phase effects in future experiments.

## 2. Material and Methods

Scripts to run the simulations and analyses, as well as the simulated data, are available in the following repository: https://doi.org/10.17863/CAM.41915

### 2.1 Simulating Experimental Data

We first created a detailed model of the phase effect (Fig. 1), based on various neural parameters (see Section 2.1.2). We then sampled this effect in a simulated experiment, based on various experimental parameters (see Section 2.1.3). For each combination of parameters, we simulated 1000 experiments in which a phasic modulation of the response variable (dichotomous or continuous) is present, and 1000 experiments in which this effect is absent. Each of the simulated experiments consisted of data from 20 virtual participants (Fig. 2A). We then applied various methods (see Section 2.2) to the simulated data and defined, for each of the combination of parameters, a method that is optimally suited (among all of the methods tested) to detect a phase effect at the group level (see Section 2.3).

#### 2.1.1. Simulating neural or behavioural responses

In each trial of a simulated experiment, a virtual response was recorded at a (e.g., neural or stimulus-specific) phase. This phase was selected in two different ways, depending on *study design* described in Section 2.1.3. A response was then determined for each trial, as follows.

For a dichotomous response variable:

If *rand*(1) >= 1-*model(phase)*, then response = “hit”, otherwise “miss”.

For a continuous response variable:

Response = *normrnd*(µ = *model(phase)*, σ = 0.7).

The terms *rand* and *normrnd* correspond to a random value drawn from a standard uniform distribution and normal distribution, respectively, and *model(phase)* corresponds to the average response (e.g., proportion of hits for a dichotomous response variable) associated with a certain phase in a detailed model of the phase effect (Fig. 1). This model was determined by the simulated neural parameters as described in the next section.

#### 2.1.2 Neural Parameters

We assigned values to the response variable at various phases (Fig. 1), which spanned a full oscillatory cycle in equidistant steps of π/96. This resulted in 192 values over a full cycle. How exactly these values varied with phase depended on following parameters:

1. *Baseline*: For all experiments, the average response (e.g., proportion of hits or scalp potential) across phase was chosen to be 0.5. Using other baselines (e.g., a value of 0) did not change the outcome of these simulations.
2. *Preferred phase*: For experiments with a phase effect present, the maximum response (most positive deflection from baseline) was assigned to a randomly chosen phase (again, with a resolution of π/96, i.e. there were 192 possibilities). The minimum response (most negative deflection from baseline) was assigned to the phase bin π away from this “preferred” phase. Note that the preferred phase was the only parameter that varied between individual subjects within each experiment.
3. *Effect size*: Effect size was defined as the area between baseline and the two (positive and negative) deflections. We chose area (rather than, for instance, the distance between positive and negative peaks) as our measure of effect size as it takes into account a larger number of phases (rather than only the most and least preferred ones). We assumed that phasic increases of a response variable (i.e. phase values yielding higher than average responses) have a matched counterpart of phase values leading to decreased responses (i.e. lower than average), as explained by VanRullen and McLelland (2013). Consequently, phase effects were constructed in such a way that the area under the positive deflection was always identical to that under the negative deflection. We tested effect sizes that correspond to a peak-to-peak modulation of between 4% and 20% (in steps of 4%) for a sinusoidal effect.
4. *Total width*: The width of a deflection (*width*_*pos*_ for the positive deflection, and *width*_*neg*_ for the negative deflection) was defined as the number of phase values covered by the deflection relative to the total number of phase values (192). In practice, *width*_*pos*_ and *width*_*neg*_ were altered by compressing the positive and negative half cycles of a sine wave, respectively, so that they covered the desired amount of phase values (Fig. 1B). The total width of the phase effect was defined as *width*_*pos*_+ *width*_*neg*_ and it could take the following values in our study: 50%, 75%, and 100%. Note that, assuming a constant effect size (area under the deflection, see *effect size* above), a reduction in total width results in an increase in peak amplitudes, as visible in Fig. 1B.
5. *Asymmetry*: Asymmetry was defined as

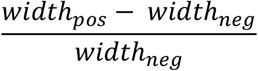

See Cole and Voytek (2019) and Schaworonkow and Nikulin (2019) for similar definitions. Note that negative or positive values of this parameter reflect a broader negative or positive deflection respectively. We simulated asymmetries between −2 and 2, in steps of 1. Again, note that, assuming a constant effect size, a change in asymmetry results in change in peak amplitudes, as visible in Fig. 1B.

#### 2.1.3. Experimental Parameters

In the context of a typical empirical study, neural parameters are not under the researcher’s control because they concern the neural mechanisms by which phase effects are reflected in neural or behavioural responses. However, researchers have control over the experimental protocol by which phase effects are measured. We therefore systematically varied the following experimental parameters:

1. *N*_*trials*_: The number of trials in total was either 192, 384, 768, or 1152. Note that the simulated data (Fig. 2A) increasingly approximates the “true” effect (Fig. 1) with increasing number of trials. 1152 trials were not tested for the continuous response variable.
2. *N*_*bins*_: For some analyses, data is sorted into phase bins in order to increase the number of trials per bin and, consequently, statistical power. Similarly, a limited number of phase bins are typically tested in studies where the phase is imposed externally (e.g., tACS) because of practical limits on the number of trials that can be tested within an experimental session. The number of phase bins simulated in our study was 4, 6, 8, 12, or 16. 16 phase bins were not tested for the continuous response variable.
3. *Study design*: We tested two common paradigms for how phases were selected for each virtual trial (see Introduction). In the first paradigm (*randomly selected*), phases were randomly selected in each trial, with 192 possible phase values as described above. In the second paradigm (*imposed externally*), only a small number of equidistant phase values was tested. In this case, the tested phase values corresponded to the centres of the phase bins used for the analysis (see *N*_*bins*_).

We note that some of the methods we tested do not divide data into phase bins (cf. Fig. 2 and Section 2.2). For these methods, we did not vary *N*_*bins*_ if phases were *randomly selected.* In the case of phases *imposed externally*, for these methods, *N*_*bins*_ only corresponds to the number of possible phases in each trial.

### 2.2 Statistical Methods

Phasic modulations of neural or behavioural responses are traditionally assessed with statistical methods that can be divided into two broad analytical categories: Parametric alignment-based methods and permutation-based methods. We here added a third category, consisting of parametric methods that are based on regression analysis and therefore avoid phase binning and re-alignment. Note that some of the permutation-based methods we investigated are also based on regression.

We conceived and evaluated 24 methods (10 parametric alignment-based, 2 parametric regression-based, and 12 permutation-based methods). Each of these is described in detail below or in the Supplementary Material. In order to keep the paper as concise and clear as possible, here we only report results for methods (1) if they have been used in previous studies or (2) if we identified them as the most sensitive (“winning”) method for at least one combination of parameters in the respective method category. Novel methods were only selected if they performed significantly better than an established method for at least one combination of parameters. These criteria led to the selection of 5 parametric alignment-based, 1 parametric regression-based, and 4 permutation-based methods, which are shown in the Results section of this article and are therefore described in detail in the following. These methods are also illustrated in Fig. 2.

#### 2.2.1 Parametric Alignment-based Methods (Fig. 2B)

All methods in this category divide data into phase bins. They only differ in their single-subject measure of the phase effect, which is calculated from the binned data and described below for each method. At the group level, all methods compare these measures against 0, here using a one-tailed, one-sample t-test. We note that several studies have compared responses across phase bins using ANOVAs (Baumgarten et al., 2015; Busch et al., 2009; Busch and VanRullen, 2010; Ng et al., 2012; Zoefel and Heil, 2013). This approach may be non-optimal since it does not take into account the hypothesized cyclical shape of the phase effect (i.e., it predicts a difference between phase bins, but, in contrast to the methods described below, not the identity of the bins that are maximally different). We therefore decided not to describe it in detail here; however, we confirmed in simulations that its sensitivity is indeed lower than that of the majority of the methods tested (data not shown).

1. MAX-OPP: The maximal response was aligned to the centre bin and the remaining bins phase-wrapped. The response of the bin opposite to the centre bin was subtracted from the average response of the two bins adjacent to the centre bin. This method is a slight modification from one that was applied by, e.g., Riecke et al. (2018).
2. MAX-ADJ: We used the same method as described in (1), but subtracted the average response of the two bins adjacent to the bin opposite to the centre bin from the average response of the two bins adjacent to the centre bin. This method was used in, e.g., Riecke et al. (2018). Note that this method cannot be used for designs with only 4 phase bins.
3. MAX-OPP VS MIN-OPP: We used the same method as described in (1) to compute difference d1. Then these steps were repeated, but after aligning individual data based on the *minimum* (instead of maximum) response, to compute difference d2. If a phase effect were present, we would expect a positive value for d1 and a negative value for d2. We therefore subtracted d2 from d1.
4. MAX-OPP-AV VS MIN-OPP-AV: We included a variation of the MAX-OPP VS MIN-OPP method for designs in which there are 8, 12, or 16 phase bins. In this case, N bins adjacent to the centre bin (e.g., if N = 2, one bin to either side of the centre bin) were used to determine d1, and M bins adjacent to and including the opposite bin (e.g., if M = 3, the one bin to either side of the opposite bin plus the opposite bin) were used to calculate d2, where

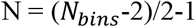

and

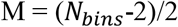 For instance, in the case of 16 phase bins, 6 phase bins adjacent to the centre bin (3 to either side) were used to calculate d1, and 6 phase bins adjacent to the bin opposite to the centre bin (3 to either side), plus this opposite bin, were used to calculate d2. For 8 phase bins, the method is very similar to its original MAX-OPP VS MIN-OPP method, with M being the only difference between the two (N=2 for both versions, but M=1 for the original version, and M=3 for the variant).
5. SINE FIT VS 0: The average response across phases was first subtracted from all phase bins (resulting in an average of 0). The response that deviated most strongly from 0 (i.e. maximum or minimum) was then aligned to the centre bin and remaining bins phase-wrapped. The sign was flipped (i.e. multiplied by −1) if the bin used for alignment corresponded to a minimum. A sine wave was then fitted to the data, excluding the centre bin and restricting the phase of the fitted sine wave so that its peak corresponded to the position of the omitted centre bin. The amplitude of the fitted sine wave was extracted. Note that this amplitude can assume negative values, corresponding to a flipped sine wave with the trough at the position of the centre bin. A similar method was used in Zoefel et al. (2018a).

#### 2.2.2 Parametric Regression-based Methods (Fig. 2C)

All the parametric regression-based methods that we tested are based on the same single-subject measure of the phase effect: Logistic (for dichotomous response variables) or linear (for continuous response variables) regression was used to test whether phase predicts responses at the single-trial level. Sine- and cosine-transformed phases were included in the regression model, yielding two circular predictors of the participant’s response. The full regression model (including phase) was then compared with an intercept-only model, using an F-Test, and yielding a p-value for each participant. The methods investigated differed in how this p-value was combined across participants. One method performed best for all parameter combinations and is therefore the only one reported here. This method, LOG/LIN REGRESS FISHER, combines the obtained p-values according to Fisher’s method (Fisher, 1950):

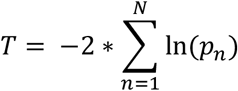

where *p*_*n*_ corresponds to the p-value for participant *n*, and T follows, under the null hypothesis, a Chi-square distribution with 2*N degrees of freedom.

A similar method has been used in Tomassini et al. (2017) and Liu and Luo (2019). However, these studies compared the two logistic regression coefficients (sine and cosine), averaged across participants, with 0 (using Hotelling’s T-squared statistic), which assumes a consistent preferred phase across participants. As this assumption does not seem to hold in all experimental scenarios (e.g., Busch et al., 2009; Riecke et al., 2018; Zoefel et al., 2018a), we randomly varied preferred phase across individuals and therefore did not include this variant of the method in our simulations.

#### 2.2.3 Permutation-based Methods (Fig. 2D)

All the permutation-based methods that we tested only differ in their single-subject measure of the phase effect, described separately for each method below. At the group level, all methods rely on a comparison of original data and a surrogate distribution. In the following, we use the term “permutation-based methods”; other terms, such as “bootstrap tests” or “resampling statistics”, are conceptually equivalent approaches (for an extensive introduction, see Good, 2005).

The common notion underlying permutation-based methods is that a hypothetical effect in the original data (e.g., a correlation between phase and an outcome measure) is abolished by randomly reassigning trial outcomes in single participants. The analysis method that was previously applied to the original dataset can then be applied to the randomized dataset which is known to lack the hypothesized effect (due to the randomization). Typically, the original data is permuted N times, thus yielding N outcomes for a given method, applied to different permutations of the same original data. These N outcomes are combined to form a distribution that is known to be produced by random fluctuations in the data, i.e. under the null hypothesis. If the original outcome is relatively unlikely to stem from the surrogate distribution, the alternative hypothesis is accepted. An advantage of this approach is that, by repeatedly “sampling” from the original data, the surrogate distribution can be used to estimate certain properties of the variable of interest (e.g., its confidence interval, cf. Schaworonkow et al., 2019). Assumptions regarding the data distribution (e.g., normality in the case of parametric methods) can therefore be avoided.

In our simulated data, the surrogate distribution was constructed by assigning single-trial responses to phase bins at random, thereby abolishing any effect of phase, and then applying the statistical methods to the permuted data. This procedure was repeated 100 times, resulting in a surrogate distribution consisting of 100 outcomes that was then compared with the single outcome obtained from the original, non-permuted data. This comparison resulted in statistical (z-) values, according to:

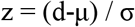

where z is the z-transformed effect size for the observed data, d is the observed data, and μ and σ are mean and standard deviation of the surrogate distribution, respectively (see VanRullen, 2016). The effect observed was considered reliable if the z-value exceeded a critical value (z = 1.645, corresponding to a significance threshold of α = 0.05, one-tailed).

For all methods tested, the single-subject measures described below were averaged across participants before the comparison with the surrogate distribution (averaged likewise in each permutation).

1. LOG/LIN REGRESS PERM: Logistic (for the dichotomous response variable) or linear (for the continuous response variable) regression with circular predictors was run as described for parametric regression-based methods. The root-mean square of the two regression coefficients (sine and cosine) was calculated. A similar method was used in Kayser et al. (2016) and McNair et al. (2019). Note that linear regression with circular predictors is mathematically equivalent to determining the amplitude of a sinusoidal wave at a given frequency with a discrete Fourier transform. Consequently, LIN REGRESS PERM is related to SINE FIT BINNED (described as method 3 below), with the only difference that the former does not require to divide data into phase bins.
2. CIRC-LIN CORR: This method was only used for the continuous response variable. Circular-linear correlation between phase and single-phase responses was calculated. This method has been used previously in Busch and VanRullen (2010) and Chakravarthi and VanRullen (2012). It is important to note that the obtained correlation coefficient is numerically equivalent to the square root of the goodness-of-fit (R^2^) from linear regression with circular predictors.
3. ITC: For the dichotomous response variable, phases were extracted separately from hit and miss trials. Phase consistency across trials (inter-trial coherence, ITC) was then calculated for each trial category, and the two ITC values summed:

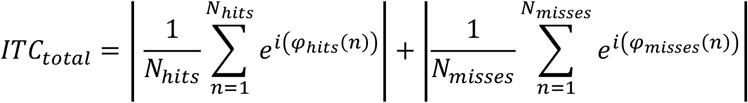

where *φ*_*hits*_(*n*) and *φ*_*misses*_(*n*) is the phase observed in trial n of all hit trials (*N*_*hits*_) and miss trials (*N*_*misses*_), respectively. For the continuous response variable, single trials were split based on the median response across trials, and *φ*_*hits*_ and *φ*_*misses*_ replaced with *φ*_*outcome*≥=*median*_ and *φ*_*outcome*≤*median*_, respectively. For dichotomous responses, this method corresponds to the Phase Opposition Sum (POS) described in detail in VanRullen (2016) and used in several studies by the same group and others (cited in the original paper). The reasoning is that under the hypothesis that the average phase for hits and misses differs, the phase consistency across trials should be higher if determined for hits and misses separately than for a mix of the two. Other variants of this method exist, all comparing phase distributions for hit and miss trials; as one of the most sensitive variants (VanRullen, 2016), POS is used here as a representative approach in this category.
4. SINE FIT BINNED: A sine wave was fitted to data divided into phase bins and averaged across trials, with phase and amplitude as free (i.e. unconstrained) parameters of this fit. The frequency of the fitted sine wave corresponded to the oscillatory frequency of interest (i.e. a single cycle in a typical experiment). Sine wave fits were achieved by running a discrete Fourier Transform and extracting the amplitude of the first non-DC output component. This method was used in Zoefel and VanRullen (2015) and Zoefel et al. (2018).

### 2.3 Evaluation of Methods

For an experiment with an effect present, if a given method yielded a significant effect at the group level (p-value < α = 0.05), this was counted as a true positive. For an experiment with effect absent, if the method yielded a significant effect, this was counted as a false positive.

The probability of a true positive (*p*_*true*_) was computed as the number of true positives observed, divided by the number of experiments with effect present (1000). Analogously, the probability of a false positive (*p*_*false*_) was computed as the number of false positives observed, divided by the number of experiments with effect absent (1000). The sensitivity of a method to detect an effect at the group level was quantified using d-prime, the standardized difference between the probabilities of true and false positives:

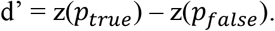

For each combination of parameters, the method yielding the highest d-prime was identified and defined as the “winning” method.

95% Confidence intervals for *p*_*true*_ and *p*_*false*_were calculated as follows:

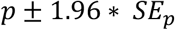

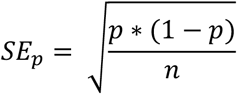

where p corresponds to *p*_*true*_ or *p*_*false*_, respectively, and n corresponds to the number of observations (i.e. experiments).

95% Confidence intervals for d-prime were calculated based on the Gourevitch and Galanter formula as follows (from Jesteadt, 2005):

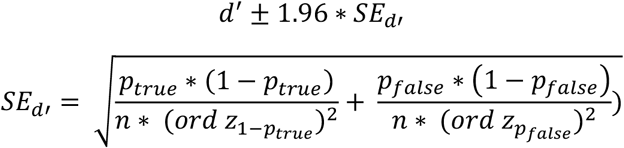

where

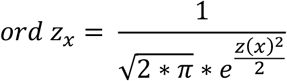

In order to determine whether two methods differed significantly in their performance, we tested whether the confidence interval of their difference included 0. A difference was defined as significant if the following condition was met:

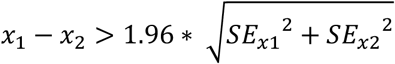

where *x*_1_ and *x*_2_ represent the measure of interest (*p*_*true*_, *p*_*false*_, or d’) from two different methods and *x*_1_ > *x*_2_.

### 2.4 Split-data Approach for Alignment-based Methods

In all of the parametric alignment-based methods tested, the phase bin used for alignment cannot be included in subsequent analyses as it is trivially an extremum. We tested whether the sensitivity of these methods can be improved if this data loss is avoided. We repeated our simulations (only for dichotomous response variable), but used a certain percentage (25%, 50%, or 75%) of the data to estimate the preferred (or “non-preferred”, for the alignment based on minimum responses) phase of each participant. This phase was then used for alignment in the remaining part of the data. Only the latter data including all phase bins was used to test for phase effects. Critically, as the phase for alignment was determined in an independent dataset, no phase bin needed to be excluded in this case. On the other hand, this procedure can reduce the number of trials used to test for phase effects (depending on the percentage of the data used to estimate preferred phase).

To apply this split-data approach, some of the methods described in Section 2.2.1 needed to be slightly modified for as described in the Supplementary Materials.

## 3. Results

We observed a very similar pattern of results for the two types of response variables. We therefore focus on results obtained for dichotomous responses, followed by a brief overview of results for continuous ones.

### 3.1 Effects of Experimental Parameters

We first pooled across all neural parameters and revealed the method with the highest sensitivity (i.e. d-prime) for each combination of experimental parameters. In Fig. 3, these highest possible sensitivities are shown for each method category. Confidence intervals for the sensitivities shown range between d’±0.12 and d’±0.16.

**Figure 3.**
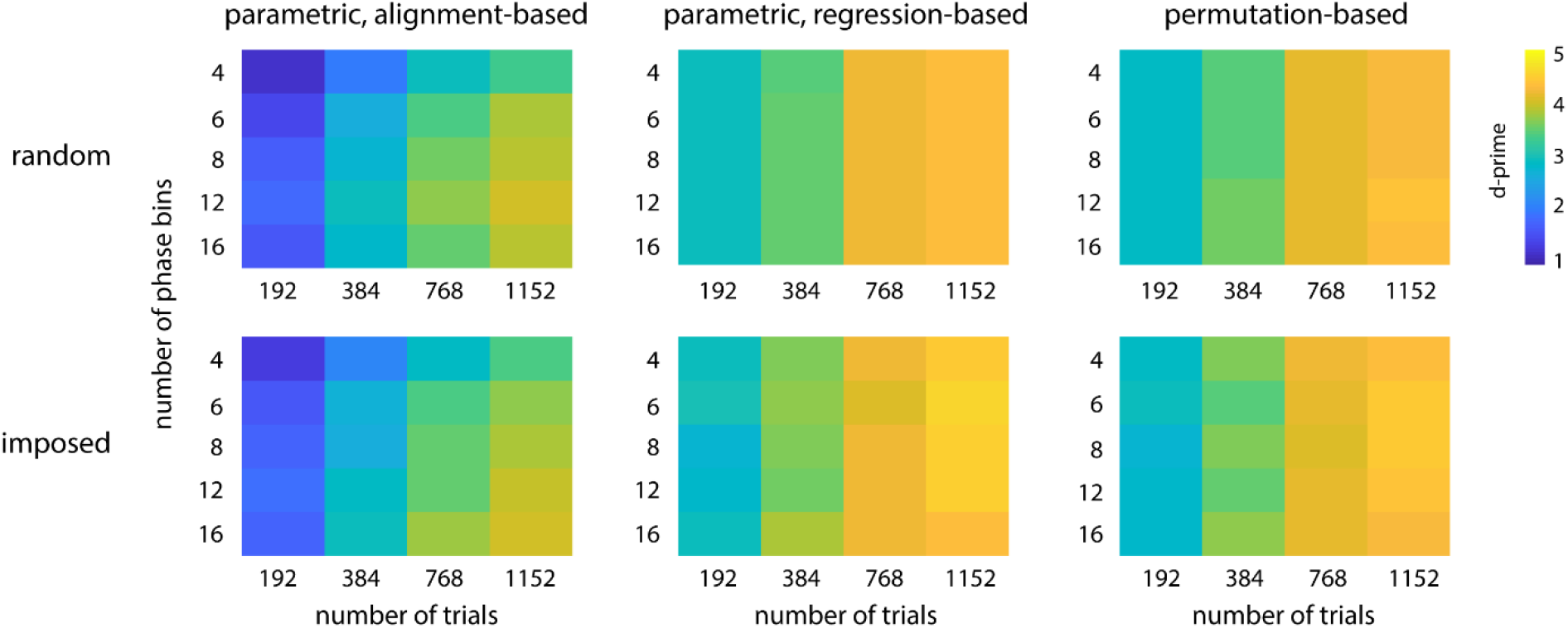
Highest possible sensitivity to detect phase effects (i.e. highest d-prime among all methods tested; color-coded) separately for all experimental parameters tested and for the three method categories. Values were averaged across all neural parameters before maximal sensitivity was determined. The confidence intervals of all sensitivities shown are in the range d’±0.12 to d’±0.16 (not shown).

We observed that parametric regression-based methods and permutation-based methods performed very similar (average d’ 3.79 vs 3.74) and generally outperformed parametric alignment-based methods (average d’ 2.89). Study design (phases *randomly selected* vs *imposed externally*) did not strongly affect the ability to detect phase effects (average d’ 3.46 vs 3.49). For all methods, sensitivity increased monotonically with *N*_*trials*_. Although less critical than *N*_*trials*_, sensitivity of parametric alignment-based methods was modulated by *N*_*bins*_, with lowest sensitivity for 4 bins, and highest sensitivity for 12 or 16 bins (depending on *N*_*trials*_ and *study design*). Most methods in the other categories do not divide data into phase bins. These methods were not affected by the number of possible phases in each trial, reflected by *N*_*bins*_ for phases *imposed externally* (see Materials and Methods). Together, these findings show that the highest possible sensitivity to detect phase effects can be achieved by using (1) regression- or permutation-based methods and (2) a high *N*_*trials*_.

Supplementary Figs. 1 and 2 show the probabilities of false and true positives, respectively, which together determine the sensitivity values shown in Fig. 3. False positive probability fluctuated around 5%, reflecting the significance threshold α applied in each simulated experiment. Consequently, none of the methods was biased towards false positives (cf. Asamoah et al., 2019), and differences in sensitivity observed were mostly driven by changes in true positive probability.

In Fig. 3, for each combination of parameters, we only show the sensitivity of a single “winning” method (that with the numerically highest value). However, in most cases, several methods performed similarly well. In Figs. 4 and 5, we therefore show sensitivity separately for different methods, together with their confidence interval. Results are shown for those experimental parameters which mostly strongly affected the identity of the best method in one or more method categories. Although not an experimental parameter, we also show results as a function of *effect size*, in order to demonstrate its positive effect on sensitivity.

**Figure 4.**
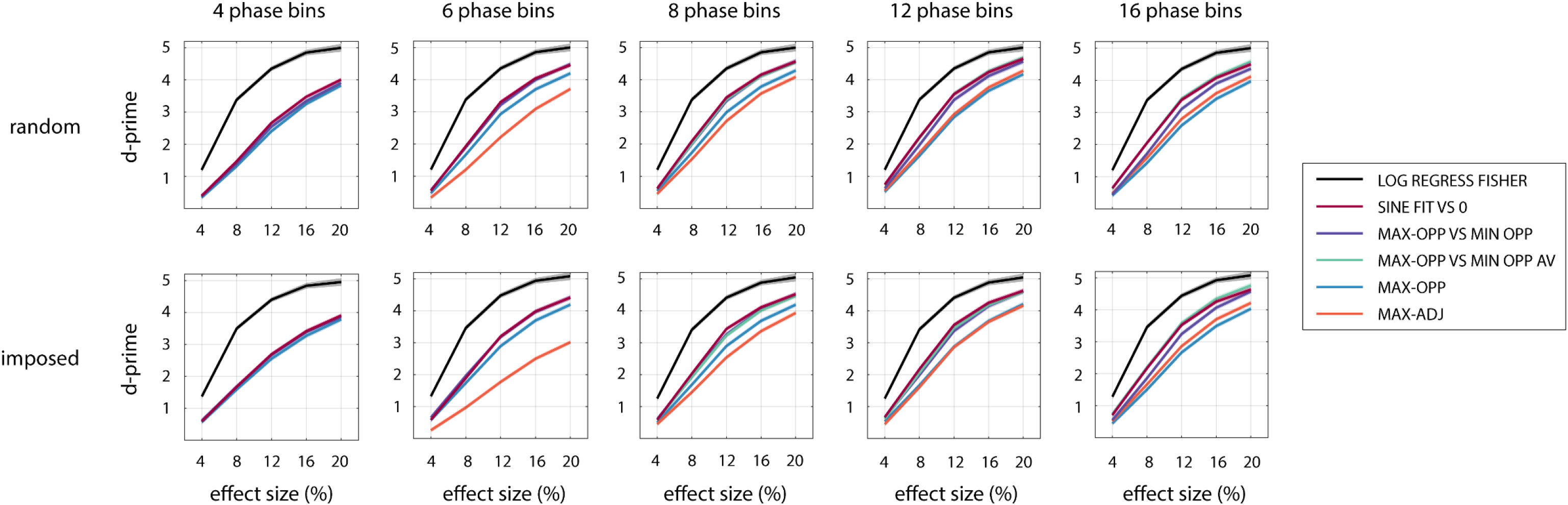
Sensitivity of the selected parametric alignment-based and parametric regression-based methods to detect phase effects, for various combinations of *N*_*bins*_, *study design*, and *effect size*. Results were averaged across *N*_*trials*_ as this parameter did not affect the identity of the winning methods. *Effect size* corresponds to a peak-to-peak modulation of performance for a sinusoidal shape (*total width* = 100%, *asymmetry =* 0). Note that MAX-ADJ is not defined for 4 phase bins, and MAX-OPP VS MIN OPP AV is not defined for 4 and 6 phase bins. Confidence intervals are shown by shaded areas – note that these are often too narrow to be visible.

**Figure 5.**
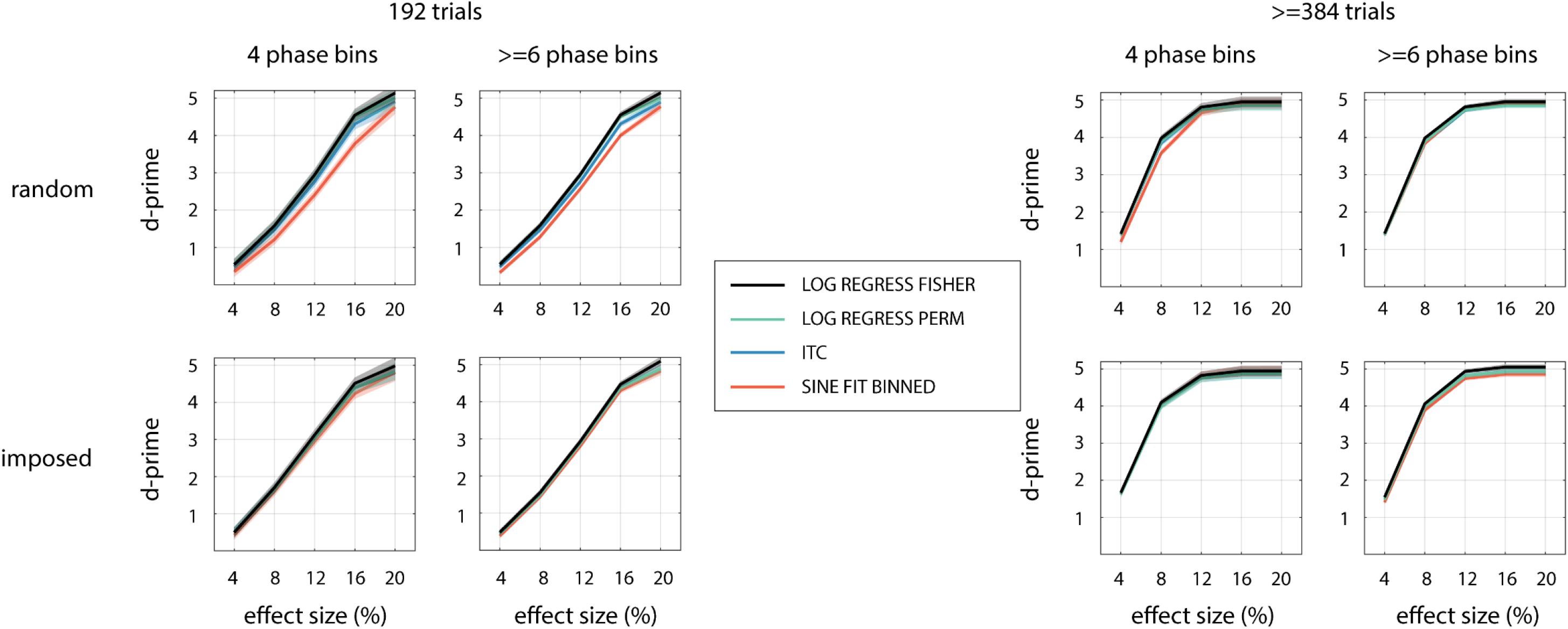
Sensitivity of the selected permutation-based and parametric regression-based methods to detect phase effects. Other conventions as in Fig. 4.

Fig. 4 shows that SINE FIT VS 0 is a parametric alignment-based method with a relatively high sensitivity in all scenarios. MAX-OPP VS MIN-OPP (for lower *N*_*bins*_) or MAX-OPP-AV VS MIN-OPP-AV (for higher *N*_*bins*_) often perform equally well. The selected parametric regression-based method (LOG REGRESS FISHER; black line) is more sensitive than all alignment-based methods and for all tested combinations of parameters.

We found similar sensitivity for LOG REGRESS FISHER and the three selected permutation-based methods (Fig. 5). Some exceptions can be seen for *randomly selected* phases, where SINE FIT BINNED is less sensitive than other methods for low *N*_*trials*_ and low *N*_*bins*_, and ITC shows a slightly reduced performance for low *N*_*trials*_. The highest sensitivity was consistently found for LOG REGRESS FISHER and LOG REGRESS PERM, which performed equally well.

Supplementary Figs. 3 and 4 show corresponding probabilities of false and true positives, respectively.

### 3.2 Effects of Neural Parameters

Next, we investigated the effect of neural parameters and their interaction with experimental parameters. In Fig. 6, we show highest possible sensitivities for different combinations of neural parameters. Confidence intervals range from d’±0.02 (for low *effect sizes*) to d’±0.12 (for high *effect sizes*).

**Figure 6.**
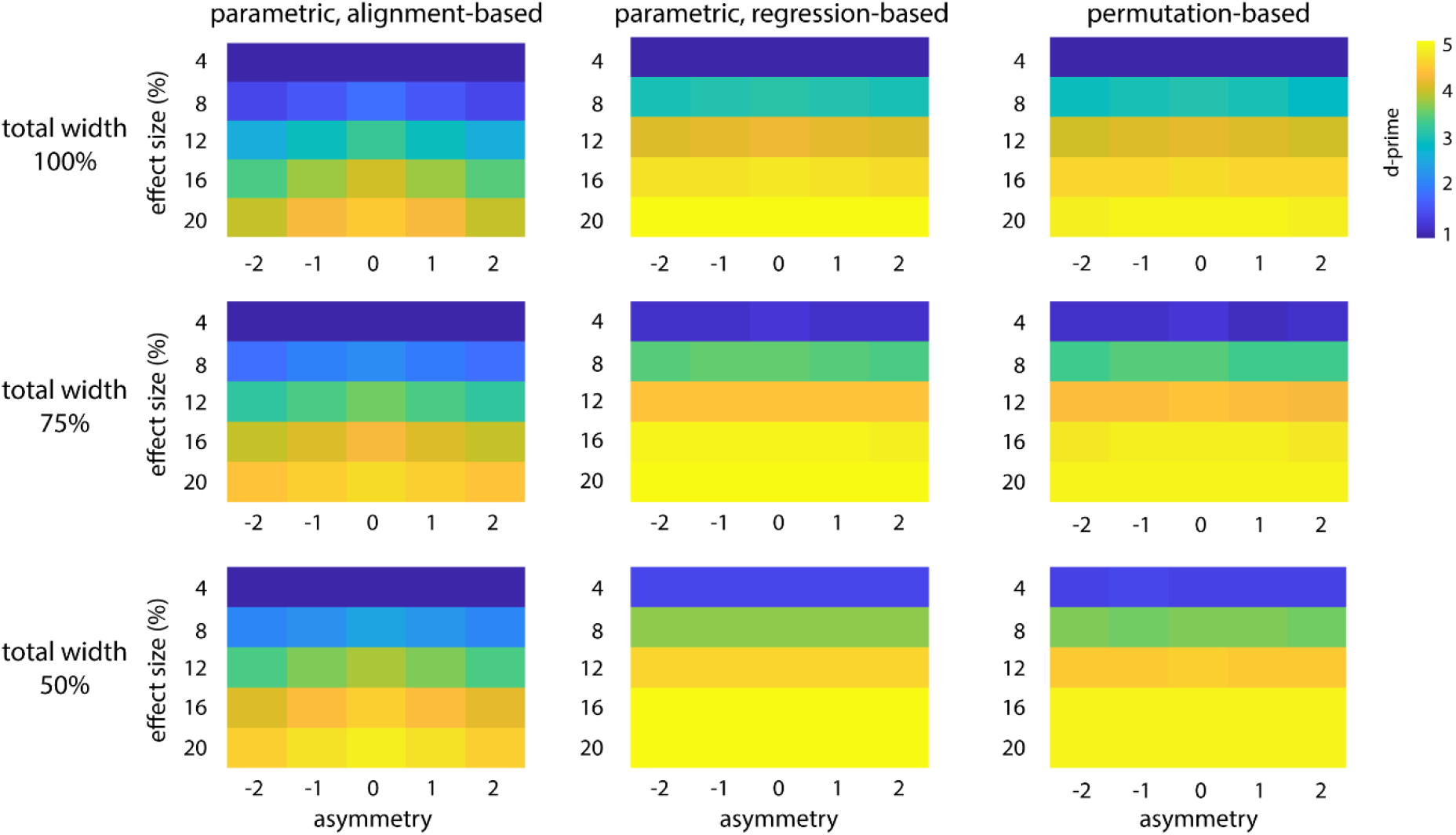
Highest possible sensitivity to detect phase effects, separately for all neural parameters and the three method categories tested. The confidence intervals of all sensitivities shown are in the range d’±0.02 (for low *effect sizes*) to d’±0.12 (for high *effect sizes*).

As already illustrated above, sensitivity to detect phase effects increased strongly with *effect size*. Sensitivity was highest for *symmetric* shapes, and this effect was most pronounced for parametric alignment-based methods (average d’ of 3.08 vs 2.73 for symmetric vs most asymmetric shape). Other method categories were affected only little by changes in asymmetry (average d’ of 3.81 vs 3.77 for parametric regression-based and 3.74 vs 3.68 for permutation-based methods, respectively). Finally, in all categories, sensitivity was modulated by *total width*, with highest sensitivity for narrow effects (increase in d’ of ~0.5 for 50% vs 100% *total width*). However, note that *effect size* was kept constant for different widths, resulting in higher peak amplitudes for narrower effects (due to *effect size* being defined based on area rather than peak amplitude; see Materials and Methods). Our results therefore suggest that the methods tested here are more effective in detecting stronger, but transient, phasic modulations than weaker modulations covering the whole oscillatory cycle.

Fig. 7 again illustrates how parametric alignment-based methods perform relative to each other. Results are shown for combinations of *N*_*bins*_ and *total width*, as the two parameters that most strongly affected the identity of the optimal method in this category. For comparison, the average sensitivity of LOG REGRESS FISHER and LOG REGRESS PERM is shown, as the two best-performing methods in the other two categories. These did not differ in their sensitivity for any of the parameter combinations shown.

**Figure 7.**
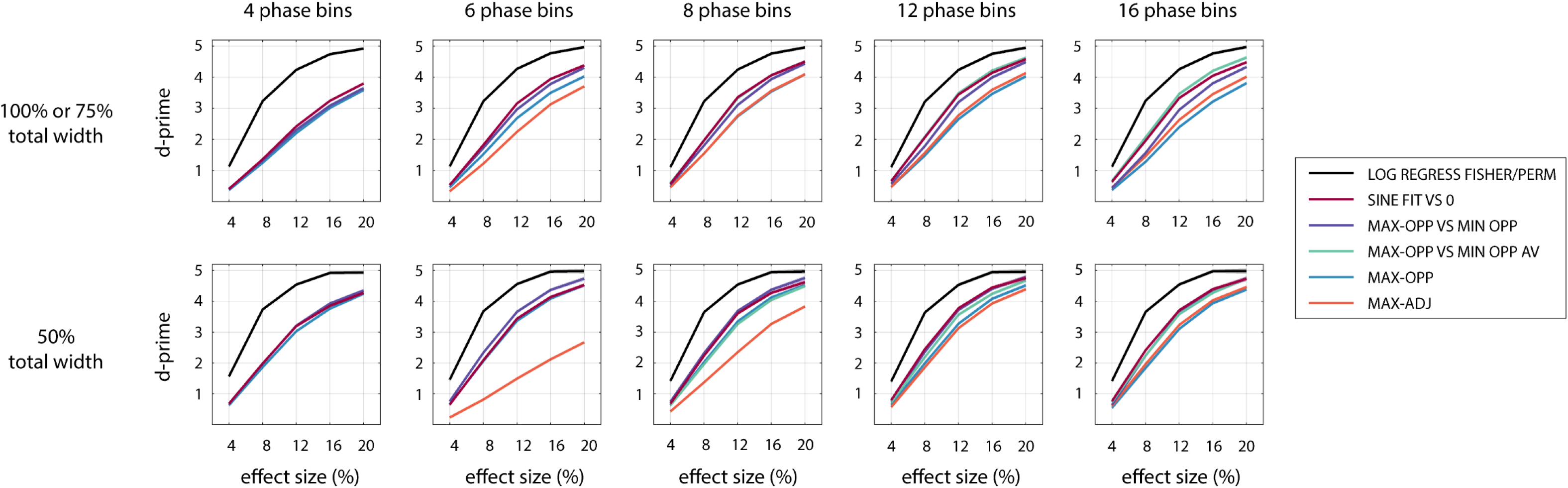
Sensitivity of the selected parametric alignment-based methods to detect phase effects, for combinations of *N*_*bins*_, *total width*, and *effect size*. The black line shows the average sensitivity for LOG REGRESS FISHER and LOG REGRESS PERM which performed equally well. For other conventions, see caption of Fig. 4.

Again, we observed that SINE FIT VS 0 is a parametric alignment-based method that is relatively sensitive in all scenarios. For narrow effects (50% *total width*), MAX-OPP VS MIN-OPP performed better for 6 and 8 phase bins. For broader effects, the sensitivity of MAX-OPP-AV VS MIN-OPP-AV increased with *N*_*bins*_, eventually becoming the winning alignment-based method for 12 or more phase bins. Regression- and permutation-based methods again outperformed alignment-based methods for all combinations of parameters.

### 3.3. Continuous Response Variable: Overview of Results

We found that the results described above can generalized to a phasic modulation of a continuous response variable. In short, (1) the best parametric regression-based (LIN REGRESS FISHER) and best permutation-based methods performed similarly and outperformed parametric alignment-based ones. (2) LIN REGRESS PERM and CIRC-LIN CORR were the permutation-based methods with the highest sensitivity for all combinations of parameters. Sensitivity of the ITC method decreased strongly, presumably because it relies on a dichotomous trial classification (“hit” vs “miss”; see Materials and Methods). (3) SINE FIT VS 0 was the most sensitive parametric alignment-based method for most parameter combinations. (4) *N*_*trials*_, *effect size*, and *effect width* (for all method categories) as well as *N*_*bins*_ and *asymmetry* (for alignment-based methods only) had a general modulatory effect on sensitivity. Fig. 8 illustrates these findings.

**Figure 8.**
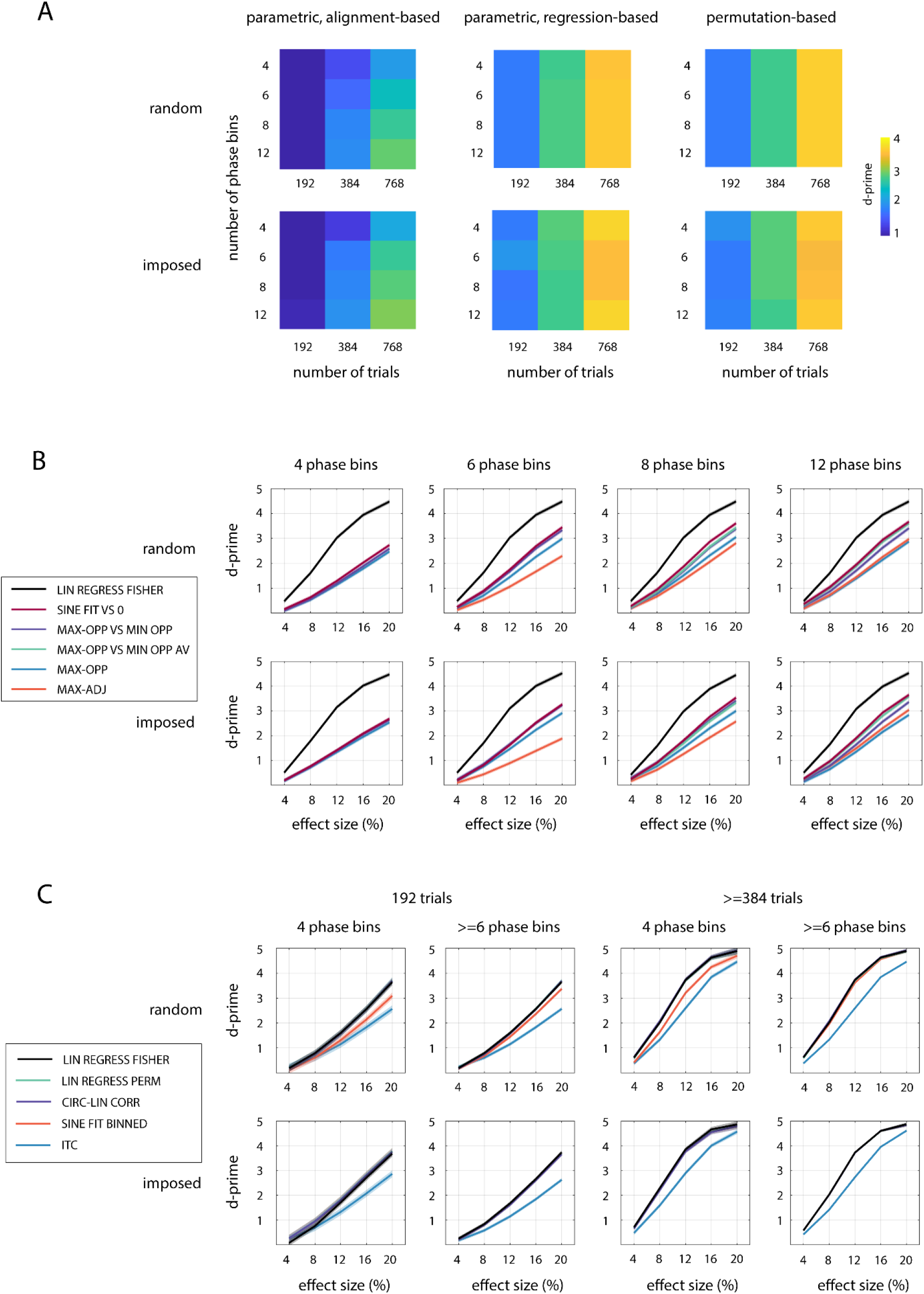
**A**. Highest possible sensitivity to detect the phasic modulation of a continuous response variable, separately for different experimental parameters. Note that overall sensitivity (across all parameters) is not comparable with that for the dichotomous response variable (Fig. 3), as it depends on the parameters that characterize the distribution from which single-trial responses are chosen (see Materials and Methods). **B,C.** Sensitivities separately for the selected parametric alignment-based (B), parametric regression-based (B,C) and permutation-based methods (C) for various simulated parameters (cf. Figs. 4,5). Note that LIN REGRESS FISHER, LIN REGRESS PERM, and CIRC-LIN CORR overlap for all parameter combinations shown in panel C. For all other conventions, see caption of Fig. 4.

### 3.4 Effectiveness of Split-data Methods for Phase Alignment

We also tested whether the sensitivity of parametric alignment-based methods can be improved by estimating individual preferred phases from a subset of data that is used exclusively for this purpose; these estimates were then used to align and test for phase effects in the remaining part of the data without the need to exclude the aligned phase bin (see Materials and Methods). Fig. 9 compares the ability of these split-data methods to detect phase effects (“split-data”) with the most common approaches described in the previous sections. We found that the split-data approach did not improve the sensitivity of alignment-based methods. That is, the availability of all trials for all but one phase bin outweighs the possibility to analyse performance in all phase bins based on a possibly smaller overall *N*_*trials*_.

**Figure 9.**
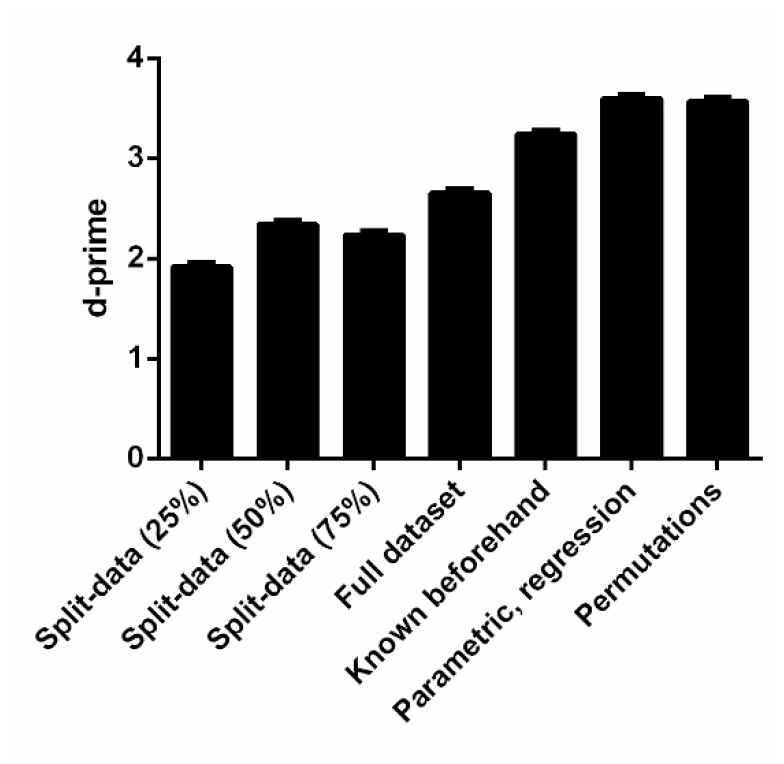
Highest possible sensitivity (i.e. highest d-prime among all methods) for split-data methods and other approaches. Results are shown for simulated experiments of 384 trials each (using other *N*_*trials*_ did not change results) and the dichotomous response variable. Sensitivity was averaged across all other parameter combinations before maximal sensitivity was determined. “Split-data”: The dataset was split, with one part used to identify the preferred phase (the percentage of data used for this purpose is indicated), the other to test for actual phase effects across all phase bins. “Full dataset”: The preferred phase was estimated in the same dataset that is used to test for phase effects, but the phase bin used for alignment was excluded from the test (i.e. the standard alignment-based approach in this paper). “Known beforehand”: The preferred phase was known beforehand (in our case, estimated in another independent dataset of 384 trials) and used to test for phase effects across all phase bins without the necessity of data exclusion. The remaining two bars show equivalent sensitivity of the winning methods in the other two categories. Error bars show the upper limits of 95% confidence intervals.

In some studies the preferred phase of individual participants may be known beforehand (e.g., from other experiments) so that phase effects can be tested based on a complete recorded dataset (without sacrificing parts of it to estimate this preferred phase). Indeed, when we assumed that this phase was known, enabling phase alignment without the necessity of excluding any data, sensitivity was improved as compared to our standard alignment-based approach. However, sensitivity was still lower than that observed for other method categories.

## 4. Discussion

There is a growing body of evidence for a rhythmic component of human perception and behaviour supported by neural oscillations (Benedetto et al., 2016; Fiebelkorn et al., 2013; Landau and Fries, 2012; VanRullen et al., 2011; Zoefel and VanRullen, 2017). Correspondingly, there is an increased interest in investigating how certain responses (from the detection of simple stimuli such as pure tones, to more complex tasks such as speech comprehension) vary with the phase of neural oscillations (e.g., Baumgarten et al., 2015; Busch et al., 2009; Henry et al., 2014; Henry and Obleser, 2012; Kayser et al., 2016; Mathewson et al., 2009; Ng et al., 2012; Riecke et al., 2018, 2015a, 2015b; Ruzzoli et al., 2019; Strauß et al., 2015; Zoefel et al., 2018a; Zoefel and Heil, 2013; Zoefel and VanRullen, 2015). However, to date, it has been unclear which methods are the most effective for detecting effects of phase on neural or behavioural responses (with the exception of a comprehensive evaluation of methods involving the comparison of two phase distributions, in VanRullen, 2016), and studies have often selected various analysis procedures on a seemingly arbitrary basis. Our goal in this work is to provide criteria to guide readers in choosing statistical procedures to be used for future studies.

### 4.1 Comparing Statistical Approaches

In all our simulations, we found that, despite being very popular in the published literature, parametric alignment-based methods have a relatively low sensitivity to detect phase effects. In contrast, we observed highest sensitivity for methods which regress single-trial responses in single participants against circular predictor variables and either combine p-values from single participants using Fisher’s method (i.e. LOG/LIN REGRESS FISHER), or compare the average regression coefficients or circular-linear correlation coefficients with a surrogate distribution (i.e. LOG/LIN REGRESS PERM or CIRC-LIN CORR). Whereas the latter approach has been used in the past (Busch and VanRullen, 2010; Chakravarthi and Vanrullen, 2012; Kayser, 2019; Kayser et al., 2016; McNair et al., 2019), as far as we know the former method has not been applied in a published manuscript (though, see Liu and Luo, 2019; Tomassini et al., 2017 for a variant of this method).

One of the most popular permutation-methods in the literature is based on testing for differences in phase consistency between hit and miss trials when analysed separately, compared to when analysed as a whole (ITC; used in, e.g., Busch et al., 2009; Dugué et al., 2011; Hanslmayr et al., 2013; Ng et al., 2012; variants of this method are summarised in VanRullen, 2016). The simulations reported here show this to be indeed among the most sensitive methods for dichotomous response variables. However, this method is not intended for, and is therefore less sensitive, to detect a phasic modulation of continuous response variables, for which we found regression-based methods to be optimal.

Although permutation-based methods performed equally well as parametric regression-based methods, they are often more computationally demanding. In addition, randomly re-assigning trials to conditions (as required for the construction of a surrogate distribution) might be problematic where this violates counterbalancing of stimulus materials. Shuffled data can include additional, item-wise variation (which is controlled in the original counter-balanced experimental data; cf. Raaijmakers et al., 1999) and which might thereby lead to invalid results. Consequently, we recommend the use of LOG/LIN REGRESS FISHER as a sensitive and efficient approach to reveal phase effects in all experiments.

### 4.2 Assumptions and Limitations

The recommendations we give here may be limited to the parameters and parameter ranges tested, and therefore may depend on the assumptions made in our study. For instance, it seems logical that enhanced responses at certain phases (relative to a baseline) have a counterpart in other, opposite phases that lead to impaired responses relative to the same baseline (VanRullen and McLelland, 2013). Based on this idea, we assumed that positive and negative effect sizes were of equal magnitude. However, the phase profile of positive and negative effects might still differ. Our neural parameters (e.g., *asymmetry*) were based on assuming that neural excitability (and therefore the shape of observed neural oscillations) directly translates into behaviour (Buzsáki and Draguhn, 2004; Lakatos et al., 2005; Peelle and Davis, 2012).

The non-sinusoidal shape of neural oscillations has increasingly been a focus of researchers’ attention (Cole and Voytek, 2017; Jones, 2016; Lozano-Soldevilla et al., 2016; Schaworonkow and Nikulin, 2019). To model the effects of non-sinusoidal waveforms on the ability to detect phase effects, we varied both asymmetry of the phase effect and the proportion of the cycle covered by the phase effect *(total width*). In this way, we simulated waveforms that closely resemble those observed in typical electrophysiological recordings. Cole and Voytek (2017) describe the human sensorimotor mu-rhythm with “one extremum (e.g., its peak) consistently sharper than the other (e.g., its trough)”. For effects covering the whole cycle (100% *total width*), our asymmetric shapes approximated this description (compare our Fig. 1B, left column, with their Fig. 1A,D). Reductions in *total width* were mostly included to address the possibility that the interaction between neural processes and externally imposed current (e.g., tACS) lead to other, potentially more complex, effect shapes. For instance, it is possible that tACS only affects performance when the applied current is at an extremum (i.e. a peak or trough).

We found that optimal parametric regression-based and permutation-based methods perform in a relatively stable manner across almost all the neural parameters tested here. Consequently, the use of these methods does not require researchers to make a-priori assumptions about the specific nature or shape of the phase effect. Nonetheless, future simulation studies should test how our preferred regression and permutation methods perform on other (e.g., “sawtooth”) waveforms that were outside the parameter set used for our simulations.

Finally, there are some plausible experimental scenarios which were not modelled in this study, for example:

1. We only simulated designs in which phases are selected randomly or controlled externally with similar or equal numbers of trials for each phase, respectively. It remains unclear how methods will perform if certain phases are tested much more often than others. This is potentially an important change as regression-based methods rely on a uniform sampling of phases.
2. We only simulated a continuous outcome variable with Gaussian distribution and it remains to be determined which other methods will be required to detect a phasic modulation of response variables with more complex distributions (e.g., with a phasic modulation that affects the shape of a distribution but not the mean).
3. The methods tested in this study employ different strategies to perform inference at the group level (e.g., comparison against chance, combining individual p-values). One common aspect across all methods applied is that single-subject measures of the phase effect can only be either 0 (if no phase effect is present, hence the null hypothesis) or larger than 0 (if a phase effect is present). This can lead to an increased number of false positives in scenarios where only a few subjects show a strong effect, while the rest has no effect. In this context, some authors have used permutation-based methods, similar to those used in our study (Gilron et al., 2017; Stelzer et al., 2013). Others have proposed to test the prevalence of the effect in the population (Allefeld et al., 2016) which, in future work, might represent an interesting alternative to the methods applied here.

### 4.3 Optimising the Detection of Neural Oscillations and their Phase

This study was designed to compare different statistical methods in their ability to detect a phasic modulation of perception, *given a reliable estimate of phase*. Nevertheless, we here briefly summarise different issues that might arise when these phases are estimated. As electrical phases are typically imposed externally and therefore known beforehand, some of these issues only concern the estimation of neural phase in electro-/neurophysiological recordings.

First, these recordings are often noisy. This will lead to uncertainty in the measurement of phase (which was not simulated here), in addition to the random fluctuations in the neural or behavioural response (which were simulated here). However, the most likely consequence of this issue is a reduction in the size of the phase effect. In our simulations, the identity of the best method was independent of effect size; thus, the best method would remain unchanged if simulations also included noise in the electrophysiological recordings. In addition, several techniques have been developed recently to reduce the impact of measurement noise (de Cheveigné and Arzounian, 2015; de Cheveigné and Parra, 2014). We also echo others in recommending the cautious use of digital filters (de Cheveigné and Nelken, 2019; Widmann et al., 2015); such filters are often necessary but can lead to spurious phase effects if applied inappropriately (Zoefel and Heil, 2013).

Second, certain variables can fluctuate over the course of an experiment in a way that can complicate reliable estimation of phase. The instantaneous frequency of alpha oscillations (~8-12 Hz) not only varies on an inter-individual level, but also within individuals (Benwell et al., 2019), and depends on factors such as luminance (Benedetto et al., 2018), cognitive load or effort (Klimesch, 2012). If these fluctuations are confined to a relatively narrow range and do not occur too rapidly, common methods, which restrict the signal to a certain frequency range before the phase is estimated (e.g., band-pass filters), should be able to capture them. More sophisticated signal processing methods exist when changes occur more rapidly, such as estimating instantaneous frequency based on peaks and troughs in the signal (Cole and Voytek, 2019). It is also plausible that the neural or behavioural response (e.g., its average across phases, termed “baseline” in this study) fluctuates over time. If these changes do not occur too rapidly (i.e. do not occur only at some of the tested phases), they should only affect the baseline but not any phasic modulation. As different baselines did not affect our results (not shown), our recommendations would remain unchanged. In addition, adaptation procedures (“staircase”) can be used to take into account fatigue or learning effects (Leek, 2001).

Third, even though the *detection* of phase effects (given a reliable phase estimate) does not seem to be strongly affected by non-sinusoidal waveforms (see previous section), the *estimation* of phase might be. Many conventional methods (e.g., Fourier Transform) assume stationary sinusoidal signals and can be biased if this assumption is not fulfilled. The approach by Cole and Voytek (2019), described above, takes this issue into account and estimates the symmetry of the extracted signal. However, it remains to be shown if this method can be applied reliably if the frequency of interest is not dominant in the recorded signal (and the estimation of peaks and troughs is difficult).

Finally, an independent question is whether the estimated phase reflects true oscillatory activity. In principle, the methods proposed here can be applied to any type of signal. The application of a narrow band-pass filter will reveal seemingly oscillatory signals at the frequency of interest, even if the data does not contain such oscillations (de Cheveigné and Nelken, 2019). Rhythmic input will necessarily lead to rhythmic brain responses, but this might merely reflect the rhythmicity of the stimulus without involving true neural oscillations (as reviewed by Zoefel et al., 2018b). This issue is increasingly recognised, and recently developed methods to identify true oscillatory activity seem promising (Doelling et al., 2019; Haller et al., 2018; Watrous et al., 2018).

### 4.4. Optimising the Detection of Phasic Modulation or Enhancement/Impairment of Behaviour

The methods presented here serve to detect a phasic *modulation* of neural or behavioural responses. If present, this modulation should reflect both in- and decreased responses relative to the average across all phases (VanRullen and McLelland, 2013). Nonetheless, when neural oscillations are perturbed by rhythmic sensory or electrical stimulation, it can be important to determine whether behaviour is enhanced, impaired, or both (but at different phases), as compared to an unstimulated baseline condition, e.g. sham stimulation (Riecke and Zoefel, 2018). It is possible that other statistical methods than those explored here will be optimal for this purpose. Future simulation studies could therefore explore statistical methods for comparison of phasic effects with non-phasic control conditions. Answering the question of whether tACS enhances behaviour relative to a control condition is of critical importance for practical applications of non-invasive brain stimulation. Only interventions that enhance desired behaviours or impair non-desired behaviours might find clinical or practical application.

We found that the most crucial experimental parameter for modulating the overall ability to detect phase effects (i.e. without necessarily influencing the identity of the best method) is the number of trials. Given this, it seems advisable to prefer simpler experimental designs with a relatively high number of trials. As evidenced by similar sensitivity for *randomly sampled* and *externally imposed* phases, determining phases post-hoc does not strongly reduce the detectability of phase effects, nor does testing only a limited number of phases (at least for methods which do not divide data into phase bins). Perhaps not surprisingly, we further observed that the most important neural parameter is the size of the hypothesized effect, which also improved sensitivity in a monotonic fashion, especially in the range of effect sizes that is commonly observed (e.g., 5-15% peak-to-peak sinusoidal modulation of behaviour). The low sensitivity of typical experiments for weak phase effects (d-prime of 1 or lower) is particularly relevant for tACS studies in which reported effect sizes are small (Gundlach et al., 2016; Neuling et al., 2012; Riecke et al., 2015a, 2015b, 2018; Riecke and Zoefel, 2018; Wilsch et al., 2018; Zoefel et al., 2018a). Transcranial brain stimulation has recently been subjected to pronounced criticism, challenging its efficacy at manipulating neural responses or behaviour (Lafon et al., 2017; Vöröslakos et al., 2018; but see Krause et al., 2019; Ruhnau et al., 2018). Our results suggest that the inconsistency of results might be partly due to the insensitivity of popular statistical methods to detect weak effects on behaviour. We anticipate two potential solutions to this problem: improving the (1) sensitivity of analysis methods (as proposed in the present paper) and/or (2) improving the efficacy of stimulation protocols (i.e. increasing neural effect sizes). It has recently been suggested that effect sizes might also be increased by adapting stimulation protocols to individual participants, as discussed elsewhere (Romei et al., 2016; Thut et al., 2017). However, these methods intrinsically depend on optimal measurement of stimulation effects on behaviour and hence are probably best built on statistical simulations like those presented here.

The pre-registration of experimental paradigms and analyses is rapidly becoming an essential step in scientific research (Nosek et al., 2018). However, the definition of detailed analysis plans in a young field of research lacking established routine procedures remains challenging (Ledgerwood, 2018). The investigation of oscillatory phase effects on behaviour is one such field in which a proliferation of analysis methods can make it difficult for researchers seeking to pre-register their analysis protocols. We hope that the simulations reported here lay the groundwork for pre-registration of tACS, EEG/MEG and other studies exploring the effect of phase on neural or behavioural responses.

## Conflict of Interest

The authors declare no competing financial interests.

## Acknowledgements

This work was supported by a grant from the German Academic Exchange Service (DAAD), the European Union’s Horizon 2020 research and innovation programme under the Marie Sklodowska-Curie grant agreement number 743482, and the Medical Research Council UK (grant number SUAG/008/RG91365). The authors thank Rufin VanRullen for helpful advice, and Nicholas Bland for providing code for the LIN REGRESS method.

## Notes

#### Summary of Updates

We tested, based on additional simulations, (1) whether results for dichotomous response variables ("hit"/"miss") can be generalized to continuous response variables and (2) how regression-based methods perform in comparison to those methods included in the original submission.

https://doi.org/10.17863/CAM.41915

