## Supplemental Material for "How to test for phasic modulation of neural and behavioural responses"

Description of additional methods tested

We tested the sensitivity of different methods to detect phase effects, and reject absent ones. For the sake of simplicity and clarity, in the main article we only report methods if they were the most sensitive method for at least one combination of simulated parameters, or if they have been used in previous research. Here we describe the remaining methods, i.e. the ones which were included in the simulations but did not reach highest sensitivity for any of the simulated parameters.

Parametric alignment-based methods

The following describes single-subject measures which are calculated from data divided into phase bins. All of these measures were subjected to a one-tailed t-test against 0 on the group level.

MAX-ABS-OPP and MAX-ABS-ADJ:

The alignment procedure described for SINE FIT VS 0 (see original paper) was used: The average response across phases was first subtracted from all phase bins (resulting in an average of 0). The response that deviated most strongly from 0 (i.e. maximum or minimum) was aligned to the centre bin and remaining bins phase-wrapped. The sign was flipped (i.e. multiplied by -1) if the bin used for alignment corresponded to a minimum. The single-subject measures described for MAX-OPP and MAX-ADJ (see original paper) were then applied to the thus aligned data. Sensitivity of MAX-ABS-OPP was higher than MAX-OPP and MAX-ADJ, but it was never the "winning" method.

MAXDIFF-ADJ:

The two phase bins with the maximal response difference of all pairs separated by a half-cycle were identified. The bin with the higher value was used for alignment. The average response of the two bins adjacent to the bin opposite to the aligned was subtracted from that of the two bins adjacent to the aligned bin.

MINVAR:

The two phase bins of all pairs separated by a half-cycle were identified for which all other bins vary minimally around the average response. The smaller response value from this pair was subtracted from the larger one.

SINE FIT VS 0 MAX:

Same as SINE FIT VS 0 (see original paper), but the maximum response (instead of maximal deviance from average) was used for alignment before the sine wave was fitted to individual data.

Parametric regression-based methods

LOG/LIN REGRESS STOUFFER:

Same as LOG/LIN REGRESS FISHER (described in Material and Methods of original paper), but Stouffer's method was used to combine p-values of individual participants:

$$T = \frac{1}{\sqrt{N}} \sum_{n=1}^{N} \phi^{-1}(1 - p_n)$$

where $p_n$ corresponds to the p-value for participant $n$, $\phi^{-1}$ represents the inverse normal cumulative distribution, and T follows, under the null hypothesis, a normal distribution.

This method was found to be similarly sensitive as LOG REGRESS FISHER.

Permutation-based methods

The following describes single-subject measures. All of these were averaged across subjects and compared with a surrogate distribution (averaged likewise). All methods described divide data into phase bins.

DIFF-NEIGHBOURS:

The average of $N_{bins}/2$ neighbouring phase bins was compared with the average of the $N_{bins}/2$ opposite phase bins. The largest difference between these two groups of phase bins was extracted from all possible comparisons. For some parameter combinations, this method was among the best permutation-based methods.

DIFF:

The difference between maximal and minimal response was used.

MAX-SURR, OPP-SURR, and MAX-OPP-SURR:

Same alignment procedure as in MAX-OPP (see original paper), followed by the extraction of the average response of the two bins adjacent to the aligned (centre) bin (MAX-SURR), the opposite bin (OPP-SURR), or the difference between the two (MAX-OPP-SURR).

MAX-OPP VS MIN-OPP SURR:

The single-subject measure described for MAX-OPP VS MIN-OPP (see original paper) was used.

SINE FIT SURR:

The amplitude of a sine wave described for SINE FIT VS 0 (see original paper) was used.

MINVAR-SURR:

The single-subject measure described for MINVAR (parametric alignment-based method, described above) was used.

Modification of methods for split-data approach

Parametric alignment-based methods (Section 2.2.1 of original paper) were modified as follows to test whether their sensitivity can be improved using a split-data approach (Section 2.4 of original paper).

1a. MAX-OPP: The response of the bin opposite to the centre bin was subtracted from that of the centre bin.

2a. MAX-ADJ: The average response of the two bins opposite to the centre bin was subtracted from that of the centre bin.

3a. MAX-OPP VS MIN-OPP: d1 and d2 were calculated by subtracting the response of the opposite bin from the centre bin.

4a. MAX-OPP VS MIN-OPP-AV: The centre bin was included in N as defined above, i.e. N = M.

5a. SINE FIT VS 0: The centre bin was included in the data used for the sine fit.

Supplementary Figures

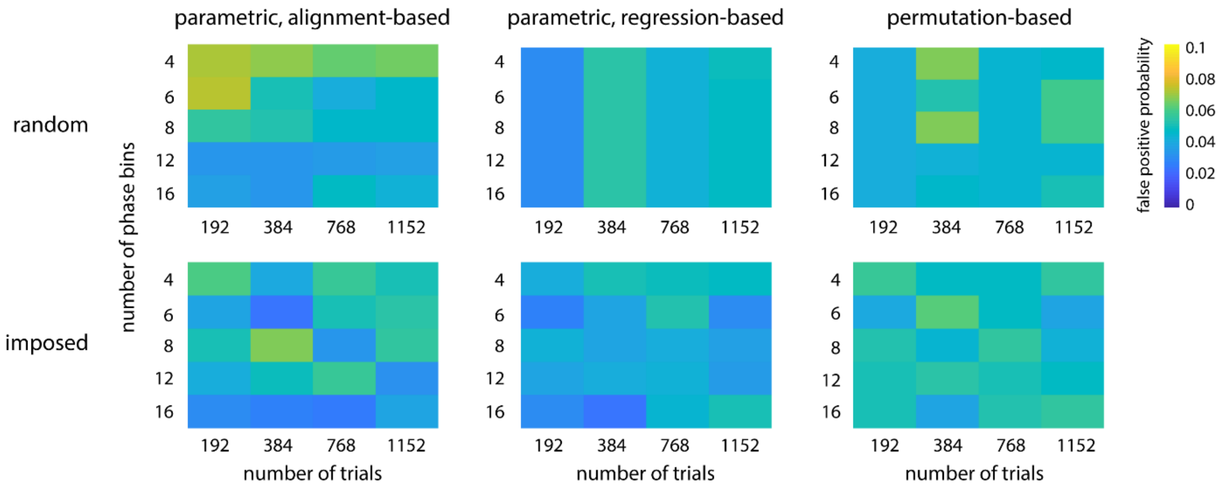


Supplementary Figure 1. Probabilities of revealing a phase effects in experiments in which these are absent (false positive). These false positive probabilities correspond to the methods with highest sensitivity (d-prime) for the combinations of experimental parameters shown. Together with the true positive probabilities depicted in Suppl. Fig. 2, they determine the sensitivities shown in Fig. 3 of the main article.

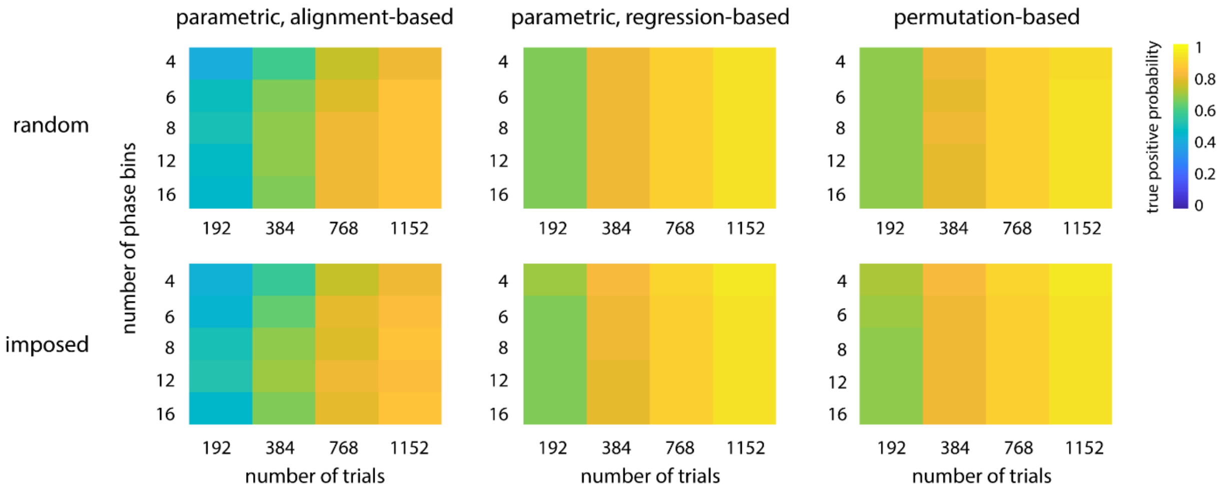


Supplementary Figure 2. Probabilities of revealing a phase effect in experiments in which this effect is present (true positive). Together with the false positive probabilities depicted in Suppl. Fig. 1, they determine the sensitivities shown in Fig. 3 of the main article.

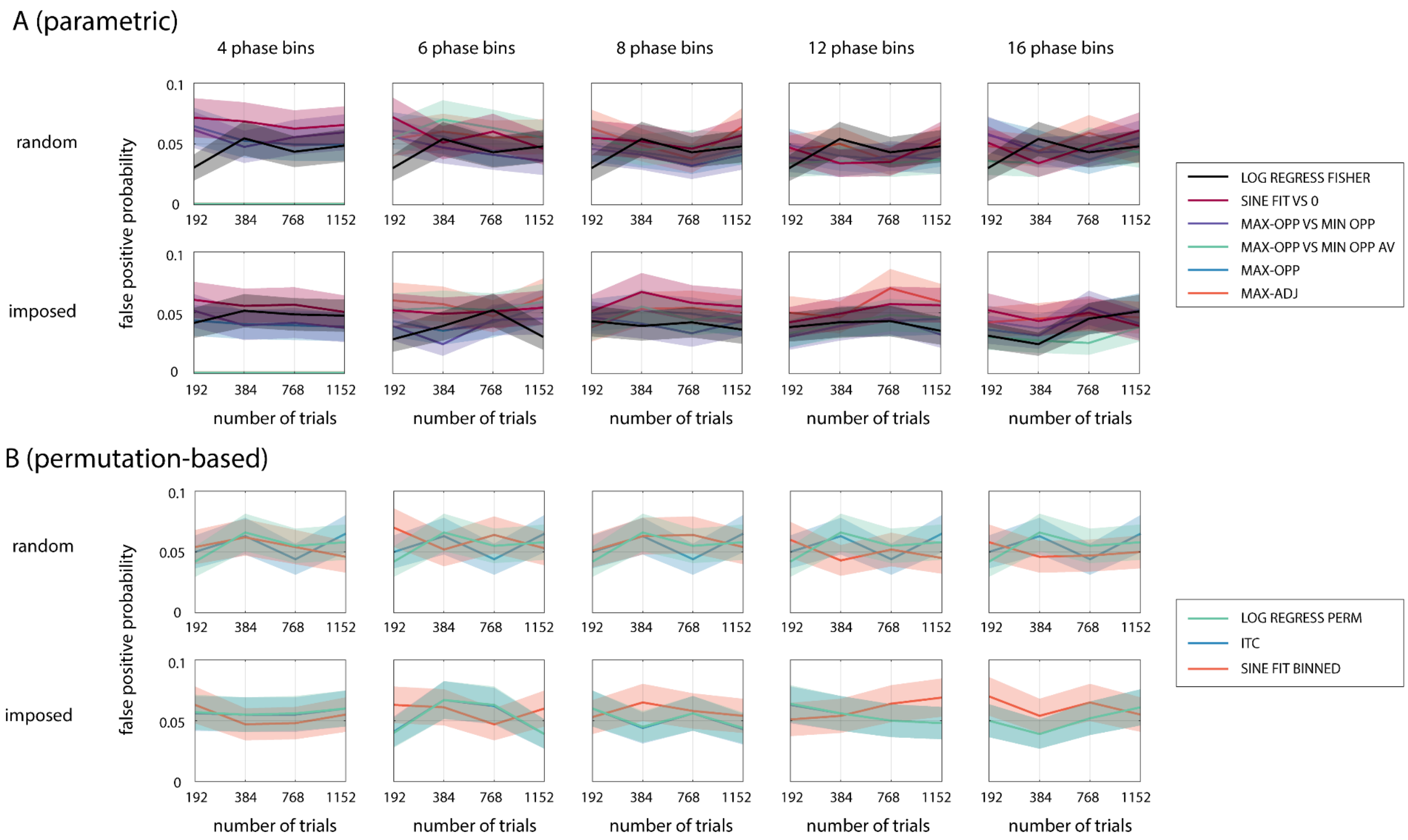


Supplementary Figure 3. Probabilities of false positives, separately for parametric (alignment- and regression-based) methods (panel A; cf. Fig. 4) and permutation-based methods (panel B; cf. Fig. 5). Confidence intervals are shown by shaded areas.

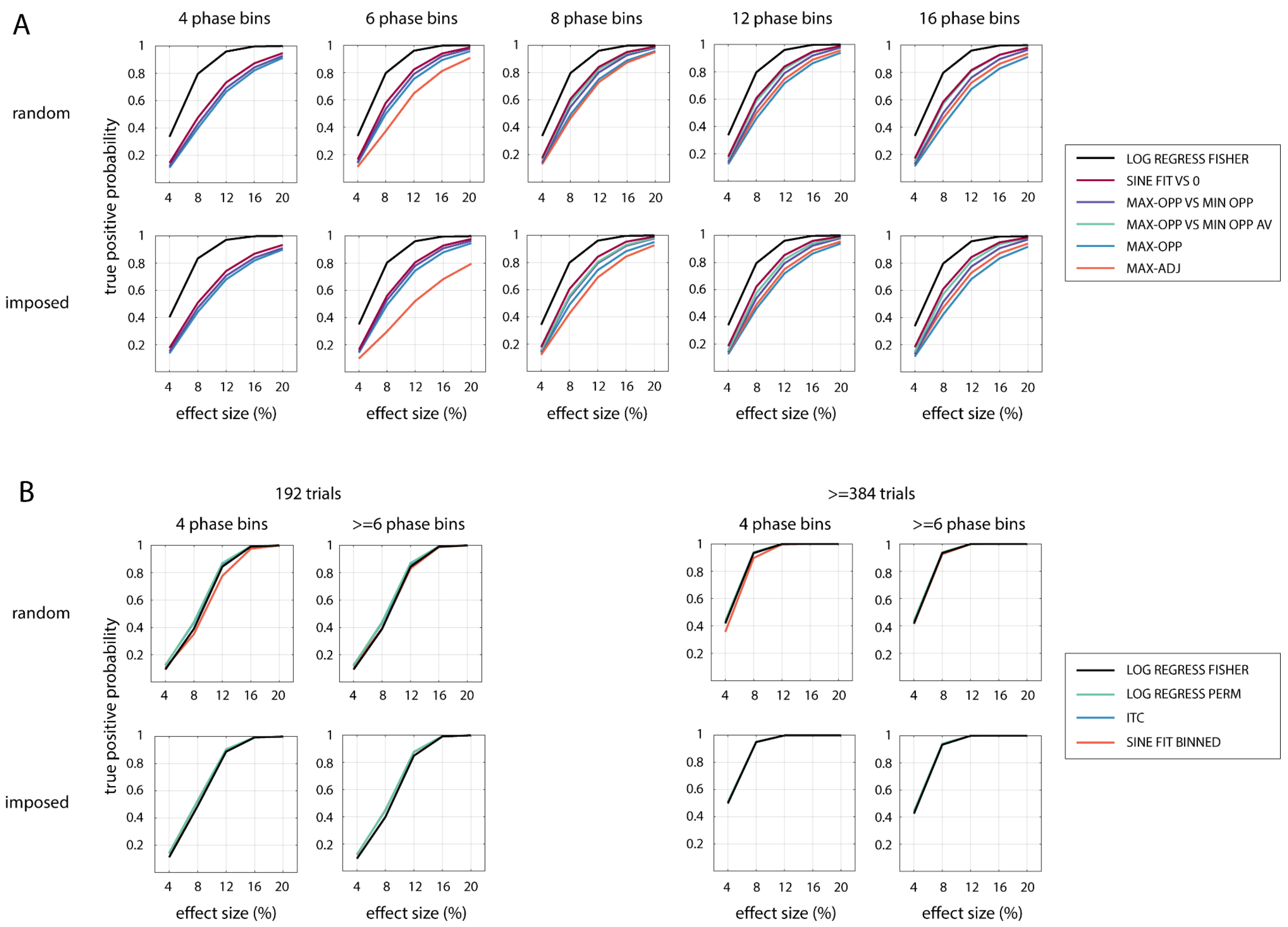
A
4 phase bins
6 phase bins
8 phase bins
12 phase bins
16 phase bins
random
imposed
true positive probability
effect size (%)
LOG REGRESS FISHER
SINE FIT VS 0
MAX-OPP VS MIN OPP
MAX-OPP VS MIN OPP AV
MAX-OPP
MAX-ADJ
B
192 trials
>=384 trials
4 phase bins
>=6 phase bins
random
imposed
true positive probability
effect size (%)
LOG REGRESS FISHER
LOG REGRESS PERM
ITC
SINE FIT BINNED

Supplementary Figure 4. Probabilities of true positives, separately for parametric alignment-based (panel A; cf. Fig. 4), parametric regression-based (panels A,B), and permutation-based methods (panel B; cf. Fig. 5). Confidence intervals are shown by shaded areas (note that these are often too narrow to be visible).
